# Tiam1-mediated synaptic plasticity drives comorbid depressive symptoms in chronic pain

**DOI:** 10.1101/2021.10.15.464592

**Authors:** Qin Ru, Yungang Lu, Ali Bin Saifullah, Francisco A Blanco, Changqun Yao, Juan P Cata, De-Pei Li, Kimberley F Tolias, Lingyong Li

## Abstract

Hyperactivity in the anterior cingulate cortex (ACC) drives comorbid depressive symptoms in chronic pain, but the cause of ACC hyperactivity is currently unclear. Ketamine, an *N*-methyl-D-aspartate receptor (NMDAR) antagonist, induces rapid and sustained antidepressant-like effects in chronic pain-induced depression in both patients and animal models. However, the mechanisms underlying ketamine’s sustained antidepressant effects remain elusive. Here, we show that Tiam1, a Rac1-specific guanine nucleotide exchange factor (GEF) that was previously identified as a critical mediator of NMDAR-dependent dendritic spine development, is activated in the ACC in chronic pain mice displaying depressive-like behaviors. Conditional deletion of Tiam1 from postnatal forebrain excitatory neurons, specific deletion of Tiam1 from ACC neurons, or pharmacological inhibition of the Tiam1-Rac1 signaling pathway prevents chronic pain-induced depressive-like behaviors in mice. Biochemical, morphological, and electrophysiological assays reveal that Tiam1 orchestrates synaptic structural and functional remodeling in ACC neurons via actin cytoskeleton reorganization and synaptic NMDAR stabilization. This Tiam1-coordinated synaptic plasticity underpins ACC hyperactivity and drives chronic pain-induced depressive-like behaviors. Ketamine induces sustained antidepressant effects in chronic pain by blocking Tiam1-mediated synaptic structural and functional plasticity in ACC neurons. Our results reveal Tiam1 as a key factor in the pathophysiology of chronic pain-induced depression and in the sustained antidepressant effects of ketamine in ACC neurons. These findings highlight Tiam1 as a potential therapeutic target for the treatment of comorbid depressive symptoms in chronic pain.

## Introduction

Chronic pain and depression are frequently comorbid, and their coexistence tends to worsen the severity of both disorders^1^. Clinical and preclinical studies have established that ACC hyperactivity is essential for driving the comorbid depressive symptoms in chronic pain^2–5^. Indeed, optogenetic activation of pyramidal neurons within the ACC is sufficient to induce depressive-like behaviors in naive mice^2^, whereas lesioning the ACC or optogenetic inhibition of pyramidal neuron hyperactivity within the ACC blocks chronic pain-induced depressive-like symptoms without affecting sensory mechanical sensitivity^2– 4,6^. While these findings argue that ACC hyperactivity underlies the comorbid depressive symptoms in chronic pain, the cause of the ACC hyperactivity remains unknown. Notably, the noncompetitive NMDAR antagonist ketamine has emerged as an effective treatment for both pain and comorbid depressive symptoms. Low, subanalgesic doses of ketamine can produce rapid, long-lasting antidepressant-like effects in patients and animal models^7,8^. However, the underlying mechanism of ketamine’s antidepressant effects has not yet been fully elucidated.

Rho GTPases (e.g. Rac1, Cdc42, and RhoA) play important roles in dendritic spine morphogenesis and synaptic plasticity by controlling the organization of the actin cytoskeleton^9^. Precise spatiotemporal regulation of Rho GTPases is mediated by activating guanine nucleotide exchange factors (GEFs) and inhibitory GTPase-activating proteins (GAPs). We previously identified the Rac1-GEF Tiam1 as a critical regulator of dendritic spine and synapse development that couples synaptic NMDAR activity to Rac1 activation and actin cytoskeletal remodeling in developing hippocampal neurons^10–12^. Here, we establish Tiam1 as a key factor in the pathophysiology of comorbid depressive symptoms in chronic pain and in the sustained antidepressant effects of ketamine through its regulation of ACC neuron maladaptive synaptic plasticity.

## Results

### Chronic pain activates Tiam1 in the ACC

The ACC appears to be a critical hub for comorbid depressive symptoms in chronic pain^2,5,6^. NMDAR-mediated enhancement of excitatory synaptic transmission in the ACC coincides with comorbid depressive symptoms in neuropathic pain^6^. Based on our previous finding that Tiam1 is a critical regulator of dendrite, spine, and synapse development that couples synaptic NMDARs to Rac1-dependent actin cytoskeletal reorganization in developing hippocampal neurons^10–12^, we explored the possible role of Tiam1-mediated synaptic remodeling in ACC hyperactivity and chronic pain-induced depression. To investigate Tiam1’s role in chronic pain-induced depressive-like symptoms, we used the spared nerve injury (SNI)-induced neuropathic pain model and the complete Freund’s adjuvant (CFA)-induced inflammatory pain model in mice (Extended Data Figs. 1a,b and 2a,b), which are both well-accepted mouse models of chronic pain^13,14^. Seven weeks after SNI surgery, mice displayed depressive/anxiety-like behaviors in several routine assays, including the forced swim test (FST) (increased immobility), the tail suspension test (TST) (increased immobility), the elevated plus maze (EPM) test (reduced open arms time), and the open field activity (OFA) test (reduced center zone time) (Extended Data Fig. 1c-f). Similar depressive/anxiety-like behaviors were observed in mice 3 weeks after CFA injection (Extended Data Fig. 2c,d), indicating that depressive/anxiety-like behaviors are reliably induced by these chronic pain models.

To investigate whether Tiam1 is activated in the ACC of chronic pain mice displaying depressive/anxiety-like behaviors, we carried out an affinity-precipitation assay to detect active Tiam1 using GST-Rac1G15A, a guanine nucleotide-free form of Rac1 that preferentially binds to activated GEFs^15^. Seven weeks after sham or SNI surgery or 3 weeks after saline or CFA injection, the active GEF assay was performed on ACC homogenates from the mice. We found that the levels of Tiam1 that precipitated with Rac1G15A from SNI or CFA mice were markedly increased compared to sham or saline controls (Fig. 1a and Extended Data Fig. 3), indicating that ACC Tiam1 is activated in chronic pain mice with depressive/anxiety-like behaviors.

**Figure 1.**
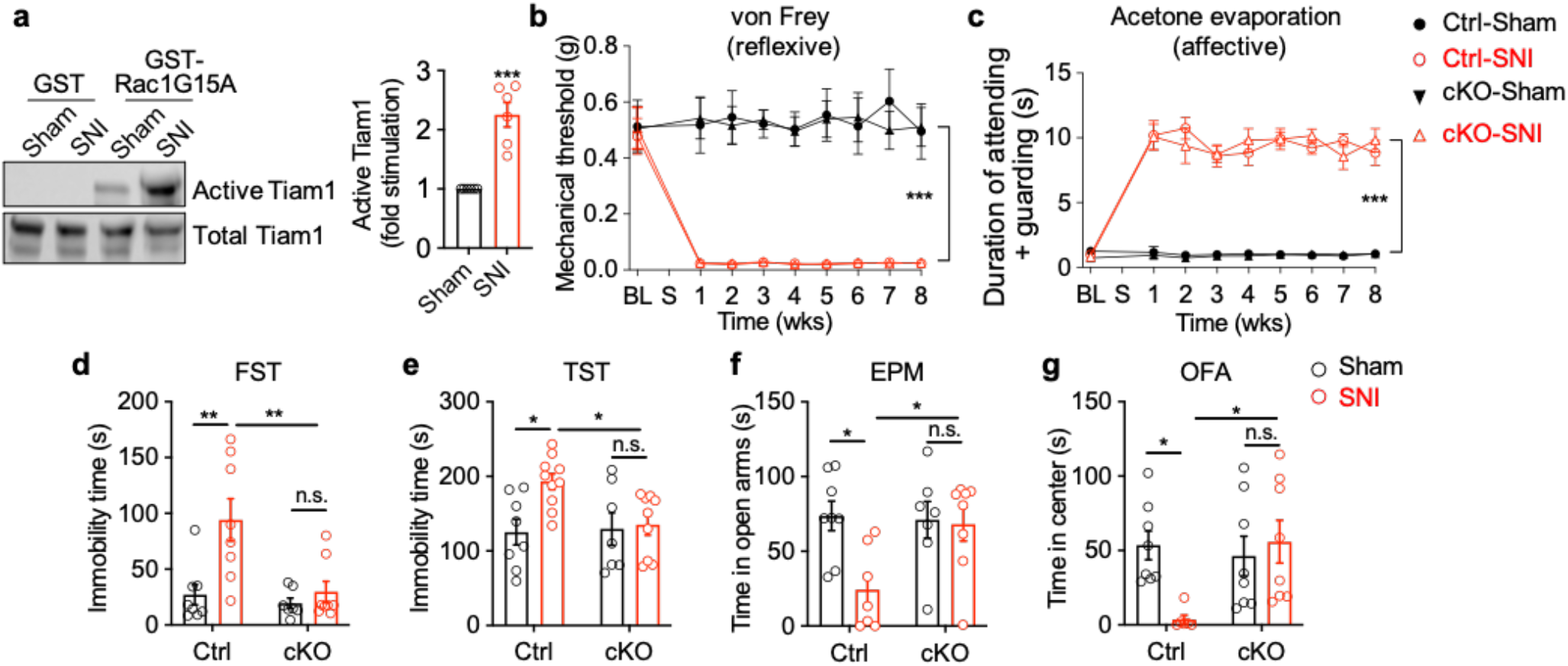
Conditional deletion of *Tiam1* from postnatal forebrain excitatory neurons prevents chronic pain-induced depressive/anxiety-like behaviors. (**a**) Representative blot and quantification of an affinity-precipitation GEF assay with GST (control) and GST-Rac1G15A (nucleotide free Rac1) showing Tiam1 activation in the ACC of mice 7 weeks after spared nerve injury (SNI) (n = 6 mice for each group. *t*_10_= 6.117, *P* < 0.0001). (**b**,**c**) Time course of SNI-induced pain reflexive behavior, demonstrated by a von Frey mechanical threshold assay (**b**, Ctrl-sham, n = 5 mice; Ctrl-SNI, n = 5 mice; cKO-sham, n = 7 mice; cKO-SNI, n = 7 mice; *F*_30,220_= 2, *P* = 0.0003) and pain affective-motivational behavior in response to acetone evaporation (**c**, n = 9 mice for each group; *F*_24,287_= 7, *P* < 0.0001) in control (Ctrl) and *Tiam1* cKO mice before and during the 8 weeks following surgery. BL: baseline, S: sham or SNI surgery. (**d-g**) Behavioral tests demonstrating that chronic neuropathic pain (7 weeks after SNI surgery) induced depressive/anxiety-like behaviors in control mice, but not in *Tiam1* cKO mice, in the forced swim test (FST) (**d**), tail suspend test (TST) (**e**), elevated plus maze (EPM) (**f**), and open field activity (OFA) (**g**) tests (Ctrl-sham, n = 8 mice; Ctrl-SNI, n = 8 mice; cKO-sham, n = 7 mice; cKO-SNI, n = 8 mice; FST, *F*_1,27_= 10.01, *P* = 0.0038; TST, *F*_1,30_= 5.486, *P* = 0.0260; EPM, *F*_1,26_= 5.673, *P* = 0.0248; OFA, *F*_1,26_= 6.3, *P* = 0.0181). Data are means ± s.e.m. * *P* < 0.05, ** *P* < 0.01, *** *P* < 0.001, n.s., no significance. Two-tailed unpaired *t*-test (a), two-way ANOVA followed by Tukey’s *post-hoc* test (b-g).

### Tiam1 in the ACC modulates chronic pain-induced depressive-like behaviors

To investigate the causal link between ACC Tiam1 activation and depressive/anxiety-like symptoms, we utilized floxed *Tiam1* mice (*Tiam1*^flox/flox^) previously generated in our lab^12^. Conditional deletion of Tiam1 from postnatal forebrain excitatory neurons (*Tiam1* cKO) was achieved by crossing *Tiam1*^flox/flox^ mice with a CaMKIIα-Cre transgenic line (Extended Data Fig. 4a). We confirmed Tiam1 loss in the forebrain via western blot analysis (Extended Data Fig. 4b). Mice lacking Tiam1 are viable, fertile, and display no gross alterations in spinal cord or brain structures^12^. *Tiam1* cKO mice also performed as well as littermate controls on the rotarod (Extended Data Fig. 4c), suggesting that they do not have deficits in motor coordination, motor learning, or balance.

Chronic pain is a complex sensory and affective experience^16^. To characterize chronic pain responses in *Tiam1* cKO mice and littermate controls subjected to sham or SNI surgery, we tested mice for both reflexive withdraw via von Frey stimuli and affective-motivational behavior in response acetone evaporation. Notably, we did not observe any difference in the mechanical withdrawal thresholds or attending and escape behaviors in control and *Tiam1* cKO mice before or during the 8 weeks following SNI surgery (Fig. 1b,c), suggesting that conditional deletion of Tiam1 from postnatal forebrain excitatory neurons does not affect peripheral nerve injury-induced pain hypersensitivity in reflexive and affective dimensions. Since *Tiam1* cKO mice develop chronic pain similar to control littermates, we next explored whether Tiam1 plays a role in chronic pain-induced depressive/anxiety-like behaviors by testing control and *Tiam1* cKO mice 7 weeks after sham or SNI surgery. We found that in contrast to SNI-treated control mice, *Tiam1* cKO mice subjected to SNI did not display depressive/anxiety-like behaviors in the FST, TST, EPM, and OFA, and instead performed similar to sham animals (Fig. 1d-g). Likewise, in the CFA-induced inflammatory pain model, *Tiam1* cKO mice displayed a marked reduction in depressive/anxiety-like behaviors compared to control mice (Extended Data Fig. 5). These data suggest that conditional deletion of Tiam1 from postnatal forebrain excitatory neurons prevents the development of chronic pain-induced depressive/anxiety-like behaviors.

To further establish the functional role of ACC Tiam1 in chronic pain-induced depression/anxiety, we specifically deleted Tiam1 from ACC neurons of *Tiam1*^flox/flox^ mice by bilateral injection of rAAV8 vector expressing Cre recombinase driven by the human Synapsin 1 promoter. An rAAV8 vector expressing GFP alone was used as the control (Fig. 2a-c). Specific deletion of Tiam1 from ACC neurons did not alter SNI-induced mechanical allodynia before or during the 8 weeks following surgery (Fig. 2d), but significantly reduced depressive-like behaviors as shown by a marked decrease in immobility times in the FST and TST (Fig. 2e,f). However, we found that Tiam1 deletion from ACC neurons had no effect on chronic pain-induced anxiety-like behaviors, as observed by no significant change in open arm time in the EPM or in center zone time in the OFA (Fig. 2g,h). These data suggest that Tiam1 expression in ACC neurons specifically modulates chronic pain-induced depressive symptoms.

**Figure 2.**
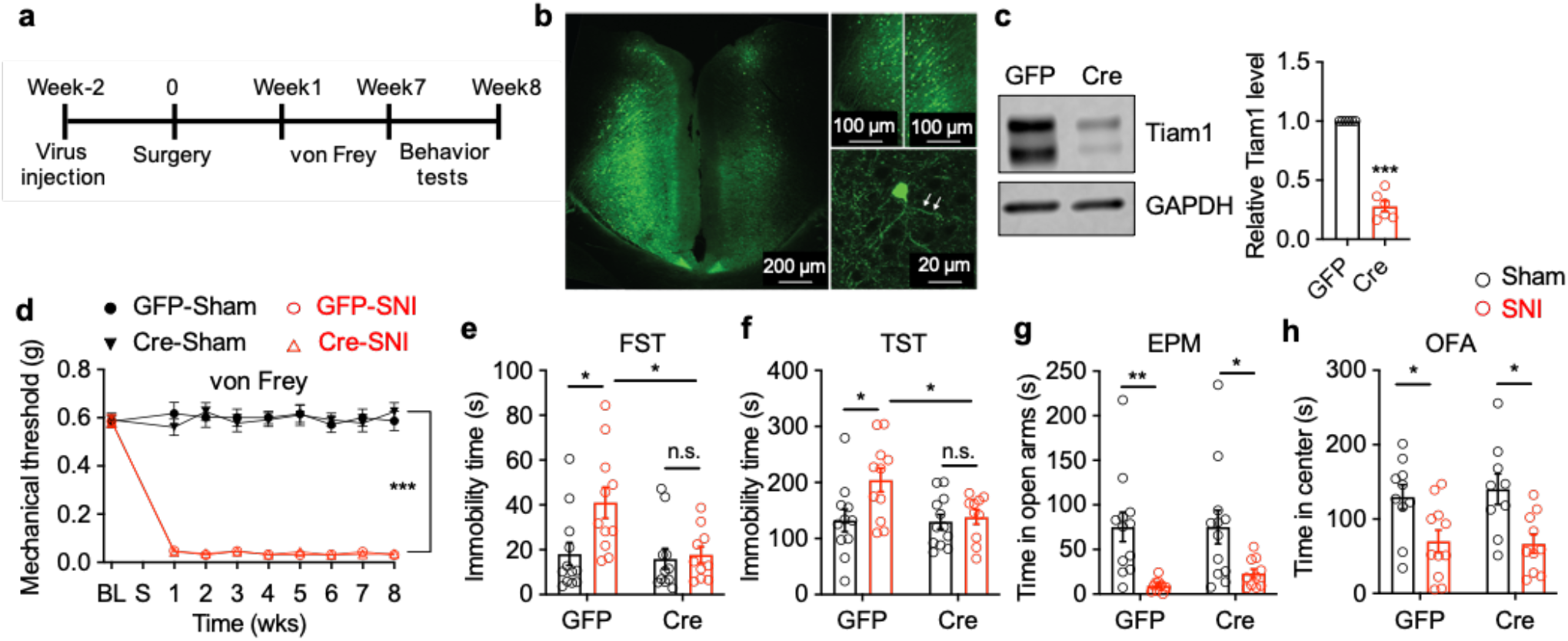
Tiam1 deletion from ACC neurons prevents chronic pain-induced depressive-like behaviors. (**a**) Experimental paradigm. (**b**) Representative picture of the ACC after bilateral injections of rAAV8 (1 μl/site) expressing Cre recombinase and/or GFP. (**c**) Western blot analysis showing Cre-mediated Tiam1 knockdown in the ACC of *Tiam1* floxed mice (n = 6 mice for each group; *t*_10_= 15.44; *P* < 0.0001). (**d**) Time course of pain hypersensitivity in response to von Frey filament stimuli in rAAV8-GFP and rAAV8-Cre-GFP injected *Tiam1* floxed mice before and during the 8 weeks following sham or SNI surgery (GFP-sham, n = 12 mice; GFP-SNI, n = 11 mice; Cre-sham, n = 11 mice; Cre-SNI, n = 11 mice; *F*_24,365_= 1, *P* < 0.0001). BL: baseline, S: sham or SNI surgery. (**e-g**) Behavioral tests demonstrating that Tiam1 deletion from ACC neurons prevented depressive-like behaviors in FST (**e**) and TST (**f**), but did not change anxiety-like behaviors in EPM (**g**) or OFA (**h**) tests in neuropathic pain mice (7 weeks after surgery) (GFP-sham, n = 12 mice; GFP-SNI, n = 11 mice; Cre-sham, n = 11 mice; Cre-SNI, n = 11 mice; FST, *F*_1,40_= 5.431, *P* = 0.0249; TST, *F*_1,40_= 5.488, *P* = 0.0242; EPM, *F*_1,40_= 0.2, *P* = 0.6258; OFA, *F*_1,36_= 0.1788, *P* = 0.6749). Data are expressed as mean ± s.e.m. * *P* < 0.05, ** *P* < 0.01, *** *P* < 0.001, n.s., no significance. Two-tailed unpaired *t*-test (c), two-way ANOVA followed by Tukey’s *post-hoc* test (d-h).

### Chronic pain induces dendritic spine remodeling and synaptic functional plasticity in ACC neurons

Basic and clinical studies have established that an underlying cause of stress-induced depression and anxiety disorders is alterations in synaptic connections in brain regions involved in mood regulation, including the prefrontal cortex, hippocampus, amygdala, and nucleus accumbens (NAc)^17,18^. Given that the ACC is a brain region that is important for the processing the emotional features of pain^2–5^, we investigated the possibility that chronic pain-induced depressive symptoms are caused by underlying synaptic alterations in the ACC. Dendritic spines are small actin-rich protrusions on dendrites that serve as the primary post-synaptic sites of excitatory synapses^19,20^. To probe for alternations in dendritic spines on ACC neurons following chronic pain, we injected a low titer of rAAV8-hSynapsin-eGFP into the ACC of mice to sparsely label ACC neurons 2 weeks before surgery. We subjected these eGFP-expressing mice to sham or SNI surgery and analyzed dendritic spines on secondary and tertiary dendrites of eGFP-positive ACC neurons 7 weeks after surgery, at a time when SNI mice display depressive-like behaviors. ACC neurons in control mice subjected to SNI showed significant increases in dendritic spine density in contrast to ACC neurons in sham control mice (Fig. 3a,b), suggesting that chronic pain does promote dendritic spinogenesis in the ACC. Since actin polymerization is a driving force for dendritic spine remodeling, we next examined whether chronic pain modulates actin dynamics in the ACC. Actin exists in two forms: monomeric globular actin (G-actin) and polymerized filamentous actin (F-actin), which is composed of aggregated G-actin. The transition between these two forms of actin is controlled by synaptic activity^20^. We used western blot analysis to measure the levels of F-actin and G-actin and found that the ratio of F-actin to G-actin, which reflects the balance between actin polymerization and depolymerization, was significantly increased in the ACC of control mice 7 weeks after SNI (Fig. 3c). Together, these data indicate that chronic pain induces synaptic structural remodeling in ACC neurons.

**Figure 3.**
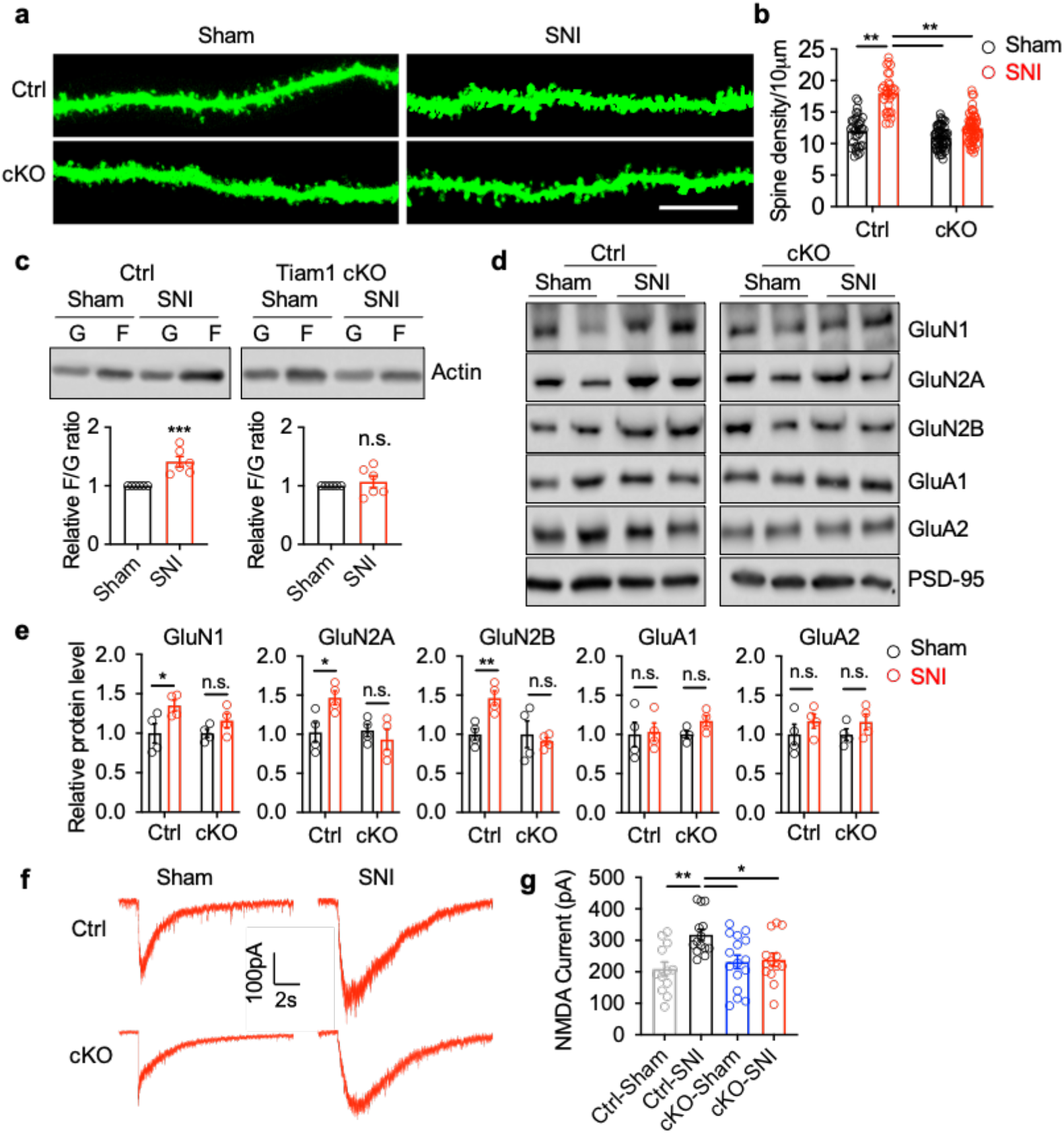
Tiam1 coordinates synaptic structural and functional plasticity of ACC neurons in chronic pain mice. (**a**,**b**) Representative confocal images (**a**) and summary of spine density (**b**) of the apical dendrites of ACC pyramidal neurons showing that chronic pain (7 weeks after SNI) increased the density of dendritic spines in control mice, but not in *Tiam1* cKO mice (Ctrl-sham, n = 26 dendrites from 3 mice; Ctrl-SNI, n = 25 dendrites from 2 mice; cKO-sham, n = 30 dendrites from 3 mice; cKO-SNI, n = 30 dendrites from 3 mice. *F*_1,180_= 41, *P* < 0.0001). Scale bar, 10 μm. (**c**) Western blot analysis revealed that the ratio of F-actin (F) to G-actin (G) in the ACC was significantly increased in control mice, but not in *Tiam1* cKO mice 7 weeks after SNI (n = 6 mice for each group. Ctrl: *t*_10_= 4.711, *P* = 0.0008; cKO: *t*_10_= 0.7141, *P* = 0.4915). (**d**,**e**) Western blot analysis revealed that in control mice (Ctrl), the levels of synaptic NMDAR subunits, but not AMPAR subunits, were significantly increased in the ACC 7 weeks after surgery (SNI). In contrast, no difference was detected in the amount of synaptic NMDAR or AMPAR subunits in the ACC of *Tiam1* cKO mice subjected to sham or SNI surgery (n = 4 mice for each group. GluN1: Ctrl, *t*_6_= 02.474, *P* = 0.0482; cKO, *t*_6_= 1.599, *P* = 0.1609. GluN2A: Ctrl, *t*_6_= 2.875, *P* = 0.0282; cKO, *t*_6_=0.7613, *P* = 0.4753. GluN2B: Ctrl, *t*_6_= 3.978, *P* = 0.0073; cKO, *t*_6_=0.4941, *P* = 0.6388. GluA1: Ctrl, *t*_6_=0.1732, *P* = 0.8682; cKO, *t*_6_= 2.143, *P* = 0.0758. GluA2: Ctrl, *t*_6_= 1.076, *P* = 0.3235; cKO, *t*_6_= 1.312, *P* = 0.2374). (**f**,**g**) Representative traces and mean changes in NMDAR currents elicited by puff application of 100 μM NMDA to ACC pyramid neurons in control and *Tiam1* cKO mice 7 weeks after sham or SNI surgery (Ctrl-sham, n = 12 neuros from 3 mice; Ctrl-SNI, n = 14 neurons from 4 mice; cKO-sham, n = 16 neurons from 4 mice; cKO-SNI, n = 13 neurons from 3 mice. *F*_3,51_= 0.4775, *P* = 0.0033). Data are expressed as mean ± s.e.m. * *P* < 0.05, ** *P* < 0.01, *** *P* < 0.001, n.s., no significance. Two-way ANOVA followed by Tukey’s *post-hoc* test (a), two-tailed unpaired *t*-test (c,e), one-way ANOVA followed by Tukey’s *post-hoc* test (g).

Synaptic structural remodeling and functional plasticity are typically highly coordinated^21–24^. For example, stress increases spine density and results in the hyperexcitability of neurons in the amygdala and NAc^25^. In the chronic pain condition, glutamate neurotransmitter system-mediated ACC hyperexcitability coincides with depressive-like consequences^6^. To determine whether chronic pain alters the glutamate receptors of ACC neurons, we isolated the postsynaptic density (PSD)-enriched membrane fraction from the ACC 7 weeks after SNI or sham surgery and measured synaptic NMDAR and AMPAR subunit protein levels with western blot analysis^26,27^. The NMDAR subunits GluN1, GluN2A, and GluN2B were increased in SNI-treated mice compared with sham control mice, whereas no differences were detected in protein expression levels of AMPAR subunits GluA1 and GluA2 (Fig. 3d,e), suggesting an increase in synaptic NMDAR levels. To measure synaptic NMDAR and AMPAR activity, we recorded NMDAR and AMPAR currents elicited by puff application of 100 μm NMDA or 200 μm AMPA directly onto the ACC pyramidal neurons. We found that chronic pain (7 weeks after SNI) markedly increased the amplitude of puff NMDA but not AMPA currents of ACC neurons (Fig. 3f,g and Extended Data Fig. 6), suggesting that chronic pain induces NMDAR-mediated hyperactivity of ACC neurons. Together, these data suggest that chronic pain-induced depressive symptoms are caused by underlying synaptic structural and functional alterations in ACC neurons.

### Tiam1 controls chronic pain-induced dendritic spine remodeling and synaptic functional plasticity

Our results suggest that chronic pain induces maladaptive synaptic plasticity in ACC neurons that is associated with chronic pain-induced depressive symptoms. Since Tiam1 in ACC neurons is required for chronic pain-induced depressive symptoms (Figs. 1,2), and Tiam1 is known to couple NMDARs to Rac1-dependent actin dynamics necessary for hippocampal spine and synapse development^10–12^, we next investigated whether chronic pain-induced depressive symptoms are determined by Tiam1-mediated dendritic spine remodeling and synaptic functional plasticity of ACC neurons. While chronic pain promotes increases in the density of dendritic spines and the F-to G-actin ratio of ACC neurons in control mice, no detectable changes in spine density or F-to G-actin ratio were observed in *Tiam1* cKO mice (Fig. 3a-c), suggesting that Tiam1 controls chronic pain-induced synaptic structural remodeling of ACC neurons. Similarly, the chronic pain-induced increases in synaptic NMDAR subunit protein levels and activity of ACC neurons observed in control mice were attenuated by the conditional deletion of Tiam1 from forebrain excitatory neurons (Fig. 3d-g), suggesting that Tiam1 regulates chronic pain-induced synaptic functional plasticity. Together, these data argue that Tiam1 coordinates synaptic structural and functional plasticity of ACC neurons, which underlies the ACC hyperactivity and the development of chronic pain-induced depression.

### Pharmacological inhibition of Tiam1 signaling alleviates chronic pain-induced depressive-like behaviors

Our genetic results indicate that Tiam1 deletion from forebrain postnatal excitatory neurons prevents the development of chronic pain-induced depressive behaviors and associated synaptic changes in the ACC. Next, we examined whether pharmacological inhibition of Tiam1 signaling could alleviate the depressive symptoms and synaptic remodeling induced by chronic pain. NSC23766 is a widely used small molecule Rac1 inhibitor that prevents Rac1 activation by the Rac1-specific GEFs Tiam1 and Trio^28^. Three-day treatment with NSC23766 (1 mg/kg, i.p.) applied 7 weeks after SNI had no effect on mechanical sensitivity threshold (Extended Data Fig. 7), but significantly alleviated SNI-induced depressive-like behaviors, as shown by marked decreased immobility in the FST and TST (Fig. 4a-c). Three-day treatment with NSC23766 7 weeks after SNI also normalized the F-to G-actin ratio, the density of dendritic spines, the synaptic NMDAR subunit levels, and the amplitude of puff NMDAR currents of ACC neurons to sham levels (Fig. 4d-j). These data suggest that pharmacological inhibition of Tiam1-Rac1 signaling alleviates chronic pain-induced depressive-like phenotypes by normalizing chronic pain-induced synaptic structural and functional remodeling of ACC neurons.

**Figure 4.**
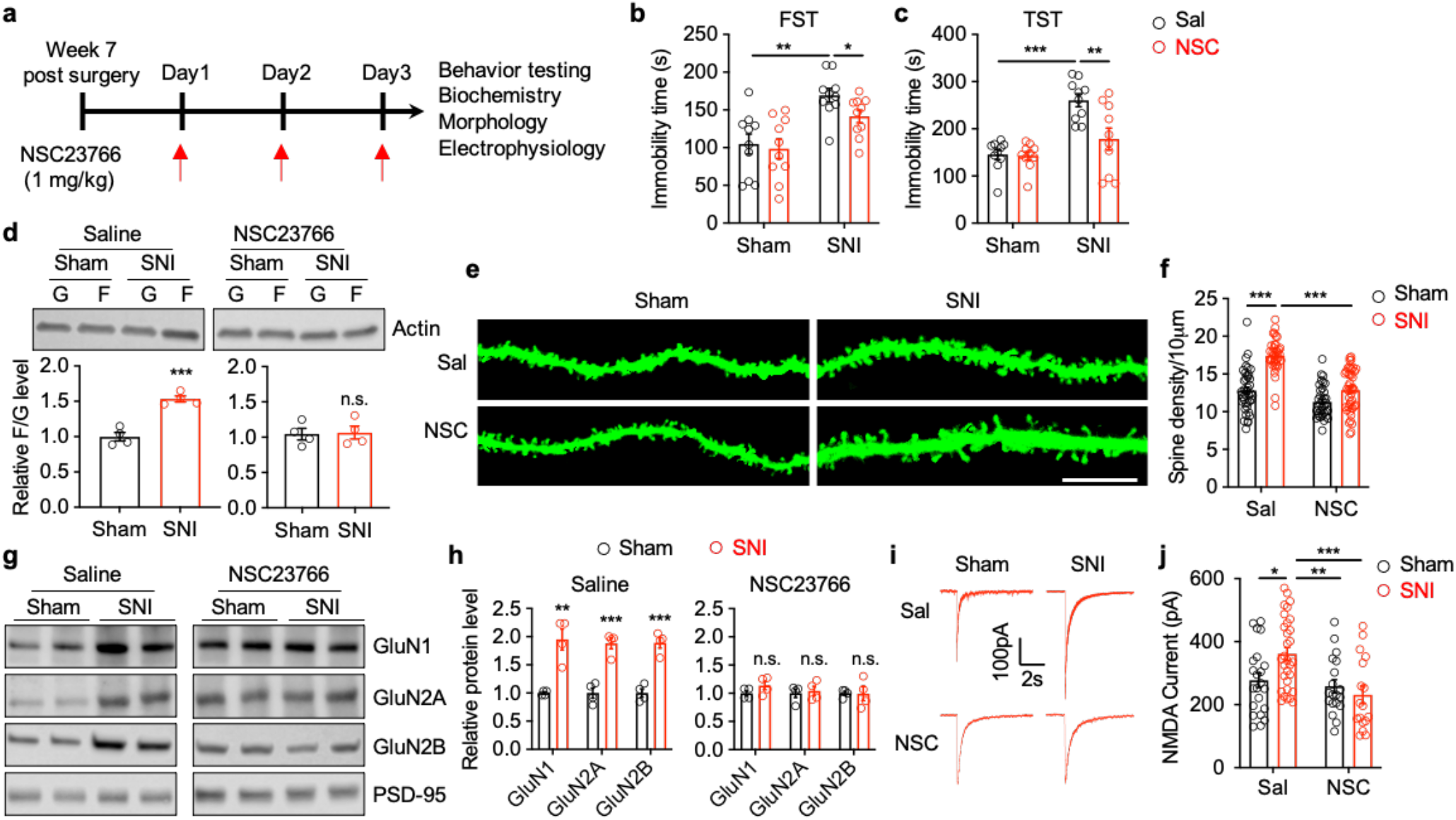
Pharmacological inhibition of Tiam1-Rac1 signaling with NSC23766 alleviates chronic pain-induced depressive-like behaviors. (**a**) Experimental paradigm. (**b**,**c**) Three-day treatment with NSC23766 7 weeks after sham or SNI surgery alleviated chronic pain-induced depressive like behaviors in the FST and TST (n = 10 mice for each group. FST, *F*_1,36_= 0.9, *P* < 0.0001; TST, *F*_1,36_= 25.11, *P* < 0.0001). (**d-j**) Three-day treatment with NSC23766 also normalized chronic pain-induced increases in the ratio of F-actin to G-actin (**d**) (n = 4 mice for each group. Saline: *t*_6_= 7.234, *P* = 0.0004; NSC23766: *t*_6_= 0.1764, *P* = 0.8657), the density of dendritic spines (**e**,**f**) (sham-saline, n = 28 dendrites from 3 mice; SNI-saline, n = 27 dendrites from 3 mice; sham-NSC23766, n = 25 dendrites from 2 mice; sham-NSC23766, n = 30 dendrites from 3 mice. *F*_1,153_= 14, *P* = 0.0002), the synaptic NMDAR subunit protein levels (**g**,**h**) (n = 4 mice for each group. Saline: GluN1, *t*_6_= 5.058, *P* = 0.0023; GluN2A, *t*_6_= 6.656, *P* = 0.0006; GluN2B, *t*_6_= 7.503, *P* = 0.0003. NSC23766: GluN1, *t*_6_= 1.546, *P* = 0.1731; GluN2A, *t*_6_= 0.3877, *P* = 0.7116; Glu2B, *t*_6_= 0.06161, *P*= 0.9529), and the NMDAR currents elicited by puff application of 100 μM NMDA to ACC pyramid neurons (**I**,**j**) (sham-saline, n = 22 neurons from 4 mice; sham-NSC23766, n = 19 neurons from 3 mice; SNI-saline, n = 31 neurons from 4 mice; SNI-NSC23766, n = 17 neurons from 3 mice. *F*_1,85_= 5.6, *P* = 0.0194). Data are means ± s.e.m. * *P* < 0.05. ** *P* < 0.01. *** *P* < 0.001. Two-way ANOVA followed by Tukey’s *post-hoc* test (b,c, f, j), two-tailed unpaired *t*-test (sham vs SNI) (d,h).

### Ketamine blocks Tiam1-dependent synaptic plasticity

The NMDAR antagonist ketamine has both analgesic and antidepressant properties^29,30^, and a single subanesthetic dose of ketamine produces rapid and sustained antidepressant-like effects in chronic pain-induced depression, without decreasing sensory hypersensitivity^7,8^. While these features have revived interest in ketamine as a treatment for comorbid depression in chronic pain, the mechanism by which ketamine mediates its effects is not fully understood. To further characterize ketamine’s effects on chronic pain-induced depression, we performed sham and SNI surgery on wild type mice. Seven weeks following surgery, we examined the time-response of a single dose of ketamine (15 mg/kg, i.p.) on mechanical sensitivity. We found that ketamine alleviated the decreased mechanical threshold observed in SNI animals 1h after administration, but this effect was no longer present at 24h post-treatment (Extended Data Fig. 8a), suggesting that the anti-allodynic effect of ketamine is transient. In contrast to its anti-allodynic effects, we found that a single injection of a subanesthetic dose of ketamine (15 mg/kg, i.p.) was sufficient to reduce neuropathic pain-induced depressive-like behaviors for at least 3 days (Fig. 5a and Extended Data Fig. 8b-e). Specifically, neuropathic pain animals showed a decrease in depressive/anxiety-like behaviors 1 h after ketamine administration in the FST (Extended Data Fig. 8b), one day after ketamine administration in the EPM test (Extended Data Fig. 8c), two days after ketamine administration in the OFA (Extended Data Fig. 8d), and three days after ketamine administration in the TST (Extended Data Fig. 8e). These data are consistent with previous reports that ketamine induces rapid and sustained antidepressant-like effects in chronic pain-induced depression^7,8^.

**Figure 5.**
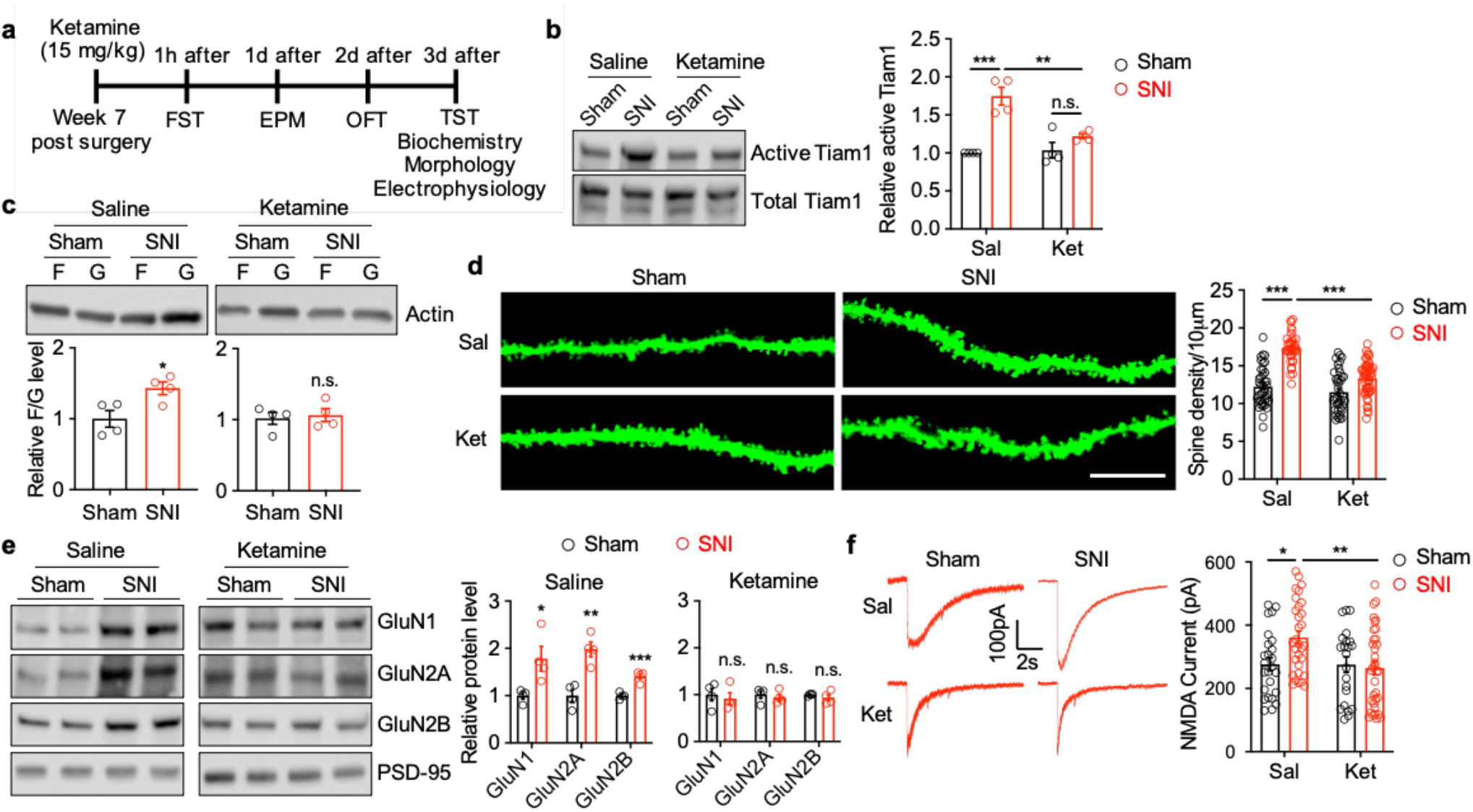
Ketamine reduces chronic pain-induced depressive symptoms by blocking Tiam1-mediated synaptic plasticity in ACC neurons. (**a**) Experimental paradigm. (**b-f**) A single subanesthetic dose of ketamine (15 mg/kg) blocked chronic pain-induced increases in the Tiam1 activity in the ACC, as demonstrated by the GST-Rac1G15A affinity-precipitation assay (**b**) (n = 4 mice for each group. *F*_1,12_= 12, *P* = 0.0040), the ratio of F-actin to G-actin in the ACC (**c**) (n = 4 mice for each group. Saline: *t*_6_= 2.938, *P* = 0.026; ketamine: *t*_6_= 0.3724, *P* = 0.7223), the density of dendritic spines in ACC neurons (**d**) (sham-saline, n = 25 dendrites from 3 mice; sham-ketamine, n = 25 dendrites from 2 mice; SNI-saline, n = 26 dendrites from 3 mice; SNI-ketamine, n = 30 dendrites from 3 mice. *F*_1,166_= 18, *P* < 0.0001), the synaptic NMDAR subunit protein levels in the ACC (**e**) (n = 4 mice for each group. Saline: GluN1, *t*_6_= 2.922, *P* = 0.0266; GluN2A, *t*_6_= 4.914, *P* = 0.0027; GluN2B, *t*_6_= 5.998, *P* = 0.001. Ketamine: GluN1, *t*_6_= 0.4536, *P* = 0.6661; GluN2A, *t*_6_= 0.5698, *P* = 0.5895; GluN2B, *t*_6_= 0.9377, *P* = 0.3846), and the NMDAR currents elicited by puff application of 100 μM NMDA to ACC pyramid neurons (**f**) (sham-saline, n = 25 neurons from 4 mice; shan-ketamine, n = 22 neurons from 3 mice; SNI-saline, n = 31 neurons from 4 mice; SNI-ketamine, n = 35 neurons from 4 mice. *F*_1,109_= 4.717, *P* = 0.0320). Data are means ± s.e.m. * *P* < 0.05. ** *P* < 0.01. *** *P* < 0.001. n.s., no significance. Two-way ANOVA followed by Tukey’s *post-hoc* test (b,d,f), two-tailed unpaired *t*-test (sham vs SNI) (c,e).

Since Tiam1 is activated in the ACC of chronic pain mice where it appears to be required for chronic pain-induced synaptic plasticity and depressive-like behaviors (Figs. 1-3), we next asked whether ketamine’s sustained antidepressant-like effects in chronic pain may be mediated, at least in part, by blocking Tiam1 function. To address this question, we first examined the effect of ketamine treatment on Tiam1 activity in the ACC of chronic pain mice. Three days after a single dose of 15 mg/kg (i.p.) ketamine or saline treatment on sham or chronic pain mice (7 weeks after SNI surgery), we performed an active GEF affinity-precipitation assay using GST-Rac1G15A on ACC homogenates. We found that ketamine treatment attenuated chronic pain-induced Tiam1 activity in the ACC (Fig. 5a,b), indicating that a single ketamine administration blocks Tiam1 activation in the ACC of SNI mice. Moreover, we found that 3 days after a single dose ketamine treatment (15 mg/kg, i.p.), the chronic pain-induced increases in the F-to G-actin ratio, the density of dendritic spines, the synaptic NMDAR subunit levels, and the amplitude of puff NMDAR currents in ACC neurons were normalized to sham uninjured mouse levels (Fig. 5c-f). These data suggest that ketamine’s sustained antidepressant-like effects in chronic pain-induced depression may be mediated by blocking Tiam1-dependent synaptic structural and functional plasticity in ACC neurons.

## Discussion

We have provided multiple lines of evidence to support the idea that Tiam1 mediates chronic pain-induced synaptic structural and functional plasticity in ACC neurons via actin cytoskeleton remodeling and synaptic NMDAR stabilization, which together determine ACC hyperactivity and depressive-like behaviors. Moreover, our results suggest that the sustained antidepressant effects of ketamine in chronic pain-induced depression are induced, at least in part, by targeting Tiam1-mediated synaptic plasticity in ACC neurons (Extended Data Fig. 9). We previously identified Tiam1 as a critical mediator of NMDAR-dependent dendritic spine development^10^. Here, in addition to showing that Tiam1 is required for chronic pain-induced spinogenesis in adult mice, we demonstrate that Tiam1 stabilizes synaptic NMDAR expression in ACC neurons, which can promote ACC hyperactivity that drives the emotional consequences of chronic pain^6^. Depressive-like behaviors induced by chronic pain often persist for weeks after the recovery from mechanical hypersensitivity^6^. Tiam1 likely contributes to the long-term nature of chronic pain-induced depressive symptoms by mediating synaptic structural and functional plasticity of ACC neurons that results in persistent modifications of ACC neuron synaptic connectivity^31,32^.

NMDARs play a pivotal role in ACC hyperactivity that drives comorbid depressive symptoms in chronic pain^6^ as well as for spinal dorsal horn hyperactivity responsible for hyperalgesia and allodynia^33,34^. As an NMDAR antagonist, ketamine’s rapid anti-nociceptive and antidepressant actions in chronic pain-induced depression (Extended Data Fig. 8) might be achieved by inhibiting NMDARs-mediated sensitization of spinal dorsal horn and ACC neurons. Notably, Tiam1 is a downstream target of NMDARs that mediates NMDAR-dependent dendritic spine morphogenesis in hippocampal neurons^10^. Our data indicate that Tiam1 is activated in the ACC in response to chronic pain (Fig. 1a and Extended Data Fig. 3) and that ketamine administration attenuates chronic pain-induced Tiam1 activation (Fig. 5b). Thus, ketamine may mediate its sustained antidepressant effects in part by inhibiting NMDAR-dependent Tiam1 activation and Tiam1-mediated maladaptive synaptic structural and functional plasticity in ACC neurons that likely underlies chronic pain-induced depression (Fig. 5 and Extended Data Fig. 9). BDNF and mTOR signaling have also been implicated in the antidepressant effects of ketamine in the models of stress-induced depression^35,36^, but whether they also contribute to ketamine’s antidepressant effects in chronic pain-induced depression and how they interact with Tiam1-medaited signaling and synaptic plasticity remains to be determined.

Besides the ACC, additional brain regions such as the amygdala, NAc, insular cortex, and prefrontal cortex are thought to be involved in the comorbidity between pain and mood disorders^37^. Interestingly, while deleting Tiam1 from ACC neurons prevented chronic pain-induced depressive-like behaviors, it had no effect on chronic pain-induced anxiety-like behaviors (Fig. 2e-h). In contrast, deletion of Tiam1 from postnatal forebrain excitatory neurons prevented both depressive- and anxiety-like behaviors in the chronic pain mice (Fig. 1d-g). Given the well documented roles of the amygdala and the NAc in mood disorders, these brain regions are strong candidates for mediating anxiety-like symptoms resulting from chronic pain^37^. Further research is needed to determine whether Tiam1 mediates synaptic plasticity in the amygdala and/or NAc driving chronic pain-induced anxiety. Taken together, this work identifies Tiam1 as a potential therapeutic target for the treatment of comorbid mood disorders in chronic pain.

## Acknowledgements

We thank Drs. Andreas Savas Tolias, Federico Scala, Surabi Veeraragavan, and K.F.T. laboratory members for technical advice and support (Baylor College of Medicine), and Dr. Yanhong Zhou for technical assistance with statistical analysis (MD Anderson Cancer Center). We also received technical assistance and resource from the BCM IDDRC Neurobehavioral Core (supported by National Institutes of Health Grant U54 HD083092). This work was supported by grants from the Department of Defense W81XWH-20-1-0790 (L.L.), and the Mission Connect/TIRR Foundation (L.L. and K.F.T.), and the National Institutes of Health NS062829 (K.F.T.).

## Author Contribution

Q.R. performed behavioral testing and morphological characterization analyses, analyzed and interpreted data; Y.L. performed the slice recordings, analyzed and interpreted data; A.B.S. performed viral injections; F.A.B. assisted with morphological characterization analyses; C.Y. assisted with animal management and behavioral testing; J.P.C assisted with ketamine experiment design; D.P.L assisted with recording experiment design and revised the article; K.F.T. designed the project, interpreted the data, and revised the article; L.L. conceived and designed the project, performed biochemical experiments and animal surgeries, analyzed and interpreted the data, drafted and edited the article.

