## Supplemental Data for "Tiam1-mediated synaptic plasticity drives comorbid depressive symptoms in chronic pain"

**This PDF file includes:**

Materials and Methods

Figs. S1 to S9

References

### Materials and Methods

**Animals.** All animal protocols were approved by the Institutional Animal Care and Use Committee of Baylor College of Medicine and were conducted in accordance with the National Institutes of Health Guidelines. For conditional knockout of Tiam1 from postnatal forebrain excitatory neurons, *Tiam1*$^{flox/flox}$ mice, generated as described (*1*), were crossed with *CaMKIIα-Cre* mice, and the resulting *Tiam1*$^{+/flox}$::*CaMKIIα-Cre* mice were then crossed with *Tiam1*$^{flox/flox}$ mice to obtain *Tiam1*$^{flox/flox}$::*CaMKIIα-Cre* (*Tiam1* cKO mice) as well as *Tiam1*$^{flox/flox}$ littermates (control) for use in experiments. Genotyping of *Tiam1* mice was determined by PCR from tail DNA using the primers: P1: ACGTGTGTTAATTAGCCAGGTTTGATGG, P2: GATCCACTAGTTCTAGAGCGGCCGAA, P3: CTACCCGGAGGAAGTGGAAGCACTACT. Genotyping of *CaMKIIα-Cre* mice was determined by PCR from tail DNA using the primers: Forward: GCATTACCGGTCGATGCAACGAGTGATGAG , Reverse: GAGTGAACGAACCTGGTCGAAATCAGTGCG. *Tiam1*$^{flox/flox}$ mice were maintained on a 129SvEv background, while *CaMKIIα-Cre* mice were maintained on a C57BL6J background. Mice were housed two to five per cage, under a 12-hour light and 12-hour dark light cycle (lights on at 06:00 and off at 18:00), with ad libitum access to food and water. All experiments used age-matched male and female mice.

**Pain models.** *Neuropathic pain.* Neuropathic pain was induced by the spared nerve injury (SNI) model (*2*). Briefly, mice were anesthetized with 2% isoflurane. A heating pad was used to maintain the core body temperature of the animals at 37 °C. An incision was made on the left lateral thigh to expose the sciatic nerve. We ligated and sectioned the common peroneal and tibial nerves (leaving the sural nerve intact) with a 5-0 silk suture under a surgical microscope. The sham procedure consisted of the same surgery without nerve ligation and section.

*Inflammatory pain.* CFA (10 μl, Sigma-Aldrich) was injected into the plantar surface of the left hind paws of mice using an insulin syringe (29G) under brief isoflurane anesthesia to induce persistent inflammatory pain. The persistence of inflammatory pain was ensured by a second CFA (10 μl) injection on the tenth day. Saline (0.9% NaCl) was injected as control.

**Viral injection.** Mice were anesthetized with 2% isoflurane and placed in a stereotaxic frame (Kopf). A heating pad was used to maintain the core body temperature of the animals at 37 °C. The coordinates were defined as dorsal-ventral (DV) from the brain surface, anterior-posterior (AP) from bregma and medio-lateral (ML) from the midline. A volume of 1μl rAAV8-hSyn-GFP or rAAV8-hSyn-Cre-GFP (UNC Vector core) was injected bilaterally in the ACC (areas 24a/24b, AP, 0.7 mm; ML, ±0.3 mm; DV, −1.5

mm) using glass micropipette attached to a Hamilton microsyringe connected to an infusion pump at a rate of 200 nl/min (*3*). After injection, the microelectrodes remained in place for 10 min and then the skin was sutured.

**Behavioral assessments of nociception.** For all behavioral tests, the experimenters performing the behavioral tests and quantitative analyses were blinded to mouse genotypes and treatments.

*Classification of mouse behaviors into reflexive and affective-motivational nociceptive responses.* The reflexive and affective-motivational nociceptive responses in mice were classified based on previous reports (*4–6*). Briefly, a cutaneous noxious stimulus can elicit several distinct behavioral responses. (i) Withdrawal reflexes: rapid reflexive withdraws of the paw that occurs in response to noxious stimulus but ceases once the stimulus is removed. (ii) Affective-motivational responses: temporally delayed (relative to the noxious stimulation), directed licking and biting of the paw (termed 'attending'), extended lifting or guarding of the paw, and/or escape responses characterized by hyperlocomotion, jumping away from the noxious stimulus, or rearing. Paw withdrawal reflexes are observed in decerebrate rodents only while the stimulus is in contact with tissue, but immediately cease once the stimulus is removed, which are classically measured in studies of hypersensitivity and involve spinal cord and brainstem circuits (*7*). In contrast, affective-motivational responses are complex behaviors, which indicate the subject's motivation and arousal to make the aversive sensations cease, by licking the affected tissue, protecting the tissue, or seeking an escape route. The affective-motivational responses require processing by limbic and cortical circuits in the brain (*8–11*).

*Mechanical reflexive assays.* To evaluate mechanical reflexive sensitivity, we used we applied a series of calibrated von Frey filaments (Stoelting, Wood Dale, IL). These filaments were applied perpendicular to the plantar surface of the hind paw with sufficient force to bend the filament. Rapid withdrawal of the paw away from the stimulus within 4 s was characterized as a positive response. If there was no response, the filament of the next greater force was applied. After a response, the filament of the next lower force was applied. We calculated the tactile stimulus force that produced a 50% likelihood of a withdrawal response using the "up-down" method (*12*, *13*).

*Thermal affective assays.* To evaluate affective-motivational responses evoked by thermal stimulation, we applied a single, unilateral 50-µl drop of acetone (evaporative cooling) to the left hind paw, and the duration of attending behavior was collected for up to 60 s after the stimulation. To prevent behavioral sensitization that can result from multiple noxious stimulations and then averaging those responses, only one drop acetone was applied on a given testing session (*5*, *6*).

**Behavioral assessments of depressive-like behaviors.** For all behavioral tests, the experimenters performing the behavioral tests and quantitative analyses were blinded to mouse genotypes and treatments. Behavioral assessments were performed during the light phase, between 09:00 and 17:00. Mice were habituated at least 1 d in the testing room before testing. On the test day, mice were transferred to the test room and were left undisturbed for at least 30 min prior to the start of testing. White noise (~60 dB) was present throughout the adaptation to the room and test.

*Rotarod.* To examine baseline motor behavior, mice were subjected to an accelerating rotarod test on two consecutive days with four trials/day. Mice rested at least 30 min between trials. The rotation speed of the rotarod increased from 4 to 40 rpm during the test. The duration of time the mice stayed on the rotarod (latency to fall) was recorded in seconds and all 8 trials were analyzed.

*Open field activity (OFA).* Mice were placed in the center of an open field arena (40 × 40 cm) and movement was recorded for 30 min with a Versamax computer assisted tracking system (Accuscan Inc., Columbus, OH). The total distance traveled was used as a measure of locomotion. The ratio between the distance traveled in a 20 × 20 cm square in the center and the total distance traveled was calculated and used as a measure of anxiety-like behavior. The area was cleaned with 75% ethanol after each test to remove olfactory cues from the apparatus.

*Elevated plus maze (EPM).* Anxiety-like behavior was tested in the EPM test for 10 minutes. Briefly, mice were placed into a maze with two 25 × 7 cm corridors with 15-cm high walls and two corridors with no walls, connected by a central square. The maze stood 50 cm above the floor. Time spent and percentages of entries into the open arms, which are measures of anxiety-like behavior, were recorded with ANY-MAZE system (Stoelting, USA). The area was cleaned with 75% ethanol after each test to remove olfactory cues from the apparatus.

*Tail suspension test (TST).* The tail suspension test was performed to study depressive-like behavior. Mice were taped by the tail to a metal bar connected to a transducer that transmitted movements to a computer. The time of immobility during a 6 min test was calculated using the ANY-MAZE system. The area was cleaned with 75% ethanol after each test to remove olfactory cues from the apparatus.

*Forced swimming test (FST).* The forced swim test was performed as described with minor modifications. In brief, individual mice were forced to swim for 6 min in a transparent plastic vessel (diameter 26 cm, height 50 cm) filled with 30 cm of water (22 ± 1°C). The immobility time was counted during a test period of 6 min using the ANY-MAZE system. The immobility time was defined as the duration a mouse floating in the water without struggling and making only small movements to keep its head above the water.

**Biochemical assays.** *Affinity-precipitation assay for Tiam1 activity.* The Tiam1 activity was measured using an affinity precipitation assay previously described (*14*). Briefly, The ACC from treated mice was isolated, homogenized in cold lysis buffer (25 mM HEPES, pH 7.4, 0.1 M NaCl, 1% NP40, 5 mM MgCl2, 10% glycerol, 1 mM DTT, 10 µg/ml leupeptin, 10 µg/ml aprotinin and 1 mM sodium orthovanadate), and centrifuged at 15,000g for 30 min. The supernatant was incubated with 30 µg of GST-Rac1G15A bound to GSH-agarose beads for 2 hr at 4 °C and mixed gently on a rocking shaker. After washing with lysis buffer for 3 times, beads were resuspended in Laemmli buffer. Samples were resolved by SDS-PAGE, transferred to nitrocellulose membrane, which was blocked with 5% fat-free milk in 0.1% Tween-PBS, and incubated with anti-Tiam1 (1:1000). Active Tiam1 was determined by Western blot analysis from the precipitated fraction and normalized to total protein (input).

*F-actin to G-action ratio.* The F-actin to G-actin ratio was determined by western blot, as previously described (*15*, *16*). Briefly, the two forms of actin differ in that F-actin is insoluble, whereas G-actin is soluble. The ACC from sham or SNI treated control and *Tiam1* cKO mice was isolated, homogenized in cold lysis buffer (10 mM K2HPO4, 100 mM NaF, 50 mM KCl, 2 mM MgCl2, 1 mM EGTA, 0.2 mM DTT, 0.5% Triton X-100, 1 mM sucrose, pH 7.0) and centrifuged at 15,000g for 30 min. Soluble actin (G-actin) was measured in the supernatant. The insoluble F-actin in the pellet was resuspended in lysis buffer plus an equal volume of buffer 2 (1.5 mM guanidine hydrochloride, 1 mM sodium acetate, 1 mM CaCl2, 1 mM ATP, 20 mM Tris-HCl, pH 7.5) and incubated on ice for 1 h to convert F-actin into soluble G-actin, with gentle mixing every 15 min. The samples were centrifuged at 15,000g for 30 min, and F-actin was measured in this supernatant. Samples from the supernatant (G-actin) and pellet (F-actin) fractions were proportionally loaded and analyzed by western blotting.

*Synaptosome preparation.* Synaptosome preparation was performed as our previous publications (*13*). The ACC from sham or SNI mice was homogenized using glass-Teflon homogenizer in 10 volumes of ice-cold HEPES-buffered sucrose (0.32 M sucrose, 1 mM EGTA, and 4 mM HEPES at pH 7.4) containing a protease inhibitor cocktail (Sigma-Aldrich). The homogenate was centrifuged at 1,000 g for 10 min at 4 °C to remove the nuclei and large debris. The supernatant was centrifuged at 10,000 g for 15 min to obtain the crude synaptosome fraction. The synaptosome pellet was lysed via hypo-osmotic shock in 9 volumes of ice-cold HEPES buffer with the protease inhibitor cocktail for 30 min. The lysate was centrifuged at 25,000 g for 20 min at 4 °C to obtain the synaptosome membrane fraction for the following immunoblotting experiments.

*Immunoblotting.* The protein samples were homogenized in RIPA buffer containing (in mM) 50 Tris-HCl (pH 7.4), 1% NP-40, 0.1%SDS, 150 NaCl, 1 EDTA, 1 Na3VO4, and 1 NaF in the presence of a proteinase inhibitor cocktail (Sigma-Aldrich). The lysates were centrifuged at 13,000 rpm for 30 min at 4

°C. The supernatant was carefully collected, and the protein concentration was measured using a DC Protein Assay Kit (Bio-Rad). A total of 30 μg of the total proteins from each sample was loaded and separated using 4−15% Tris-HCl SDS-PAGE gels. The resolved proteins were transferred to an Immobilon-P membrane (Millipore). The membrane was treated with 5% nonfat dry milk in TBST at 25 °C for 1 hr and then incubated in TBST supplemented with 0.1% Triton X-100 and 1% BSA and primary antibodies overnight at 4 °C. The membrane was washed three times and then incubated with horseradish peroxidase-conjugated secondary antibodies for 1 h at 25 °C. The protein band was revealed using an ECL Plus Detection Kit (Thermo Fisher Scientific, Waltham, MA), and the protein band density was quantified with the Odyssey Fc Imager (LI-COR Biosciences, Lincoln, NE) and normalized to the control protein band on the same blot. Tiam1 was detected using rabbit anti-Tiam1 antibody (sc-872, 1:1,000; Santa Cruz); Actin was detected using mouse anti-Actin antibody (MAB1501, 1:10,000; Millipore); GluN1 was detected using rabbit anti-GluN1 antibody (G8913, 1:1,000; Sigma); GluN2A was detected using rabbit anti-GluN2A (PA5-35377, 1:1,000; Thermo Fisher Scientific); GluN2B was detected using anti-mouse GluN2B (75-002, 1:1,000; NeuroMab); GluA1 was detected using mouse anti-GluA1 antibody (75-327, 1:1,000; NeuroMab); GluA2 was detected by using rabbit anti-GluA2 antibody (ab10529, 1:1,000; Millipore); PSD-95 was detected by using rabbit anti-PSD-95 antibody (ab18258, 1:2,000; abcam); GAPDH was detected by using mouse anti-GAPDH (sc-47724, 1:1,000; Santa Cruz).

**Morphological analysis.** To observe the effects of Tiam1 on spine remodeling in the ACC, rAAV8-hSyn-eGFP (UNC vector core) was injected bilaterally in the ACC and was used to specifically label the neurons. ACC sections (40 μm thick) were collected from mice perfused with 4% PFA, and only dendritic spines on neurons labelled with eGFP were selected for the spine analysis in a blinded manner as previously described(*1*). All spine images were captured using a Laser Scanning Confocal Microscope (LSCM, Zeiss LSM 880, Germany) with a 63x oil 210 immersion objective. Z series were taken at an interval of 0.37 μm for each dendrite. Spine morphometric analysis was done in a blinded manner using Imaris software (Bitplane Scientific Software) as previously described (*17*).

**Brain slice preparation and electrophysiology.** The brain slices containing the ACC were obtained following previously protocol (*18*) with some modification. In brief, mice were anesthetized with 3% isoflurane and decapitated. Their brains were rapidly removed and collected into ice-cold (~ 0 °C) oxygenated NMDG (N-methyl-d-glutamine) solution containing 93 mM NMDG, 93 mM HCl, 2.5 mM KCl, 1.2 mM $NaH_2PO_4$, 30 mM $NaHCO_3$, 20 mM HEPES, 25 mM glucose, 5 mM sodium ascorbate, 2 mM thiourea, 3 mM sodium pyruvate, 10 mM $MgSO_4$ and 0.5 mM $CaCl_2$, pH 7.35 (all from Sigma-

Aldrich). Coronal slices were cut 300-μm-thick using a Leica VT1200 microtome following coordinates provided in the Allen Brain Atlas for adult mice (http://atlas.brain-map.org). The slices were subsequently incubated at 34.0 ± 0.5 °C in oxygenated NMDG solution for 10-15 min before being transferred to the artificial cerebrospinal fluid (ACSF) solution containing: 125 mM NaCl, 2.5 mM KCl, 1.25 mM $NaH_2PO_4$, 25 mM $NaHCO_3$, 1 mM MgCl2, 11.1 mM glucose and 2 mM $CaCl_2$, pH 7.4 (all from Sigma-Aldrich) for about half hour. The slices were allowed to recover in ACSF equilibrated with bubbling with 95%O2/5%CO2 gas mixture at room temperature (approximately 25 °C) for at least 1 h before experiments. During the recordings, individual slices were transferred to a customized recording chamber and submerged in the chamber continuously perfused with oxygenated ACSF warmed to 32–34 °C by passing it through a feedback-controlled in-line heater (Temperature controller VII, Luigs & Neumann GmbH). Recorded cells were generally located 15–60 μm deep under the slice surface.

The pipette internal solution contained (in mM) 135.0 potassium gluconate, 5.0 TEA, 2.0 $MgCl_2$, 0.5 $CaCl_2$, 5.0 HEPES, 5.0 EGTA, 5.0 Mg-ATP, 0.5 Na-GTP and 10 lidocaine (lignocaine) N-ethyl bromide (adjusted to pH 7.2–7.4 with 1 M KOH; 290–300 mOsmol/L). NMDAR- or AMPAR-mediated currents were elicited by puff application of 100 μM NMDA or 200 μM AMPA to the recorded neuron at a holding potential of -60 mV. Positive pressure (4 p.s.i., 15 ms; Picospritzer III) was applied, and puff application of the vehicle produced no currents. The tip of the puff pipette was placed 100~150 μm away from the recorded neurons in the presence of 1 μM TTX. To minimize the $Mg^{2+}$ block of NMDARs, the puff NMDA currents were recorded in an extracellular solution containing no $Mg^{2+}$ and 10 μM glycine (*13*, *19*).

**Statistical analyses.** All statistical analyses were performed using Prism 9 software (GraphPad Software Inc., San Diego, CA). No statistical methods were used to pre-determine sample sizes but our sample sizes are similar to those reported in previous publications (*13*, *20*). The normality test was performed by the Shapiro–Wilk test. Data that met these two conditions were analyzed using a two-tailed unpaired or paired t-test, one-factor analysis of variance (ANOVA) and repeated-measures ANOVA followed by Tukey's multiple comparisons test. Data are presented as means ± s.e.m. All behavioral, electrophysiological, biochemical, and morphological data were obtained by counterbalancing experimental conditions with controls. Statistical significance was accepted when $P < 0.05$.

### Figs. S1 to S9

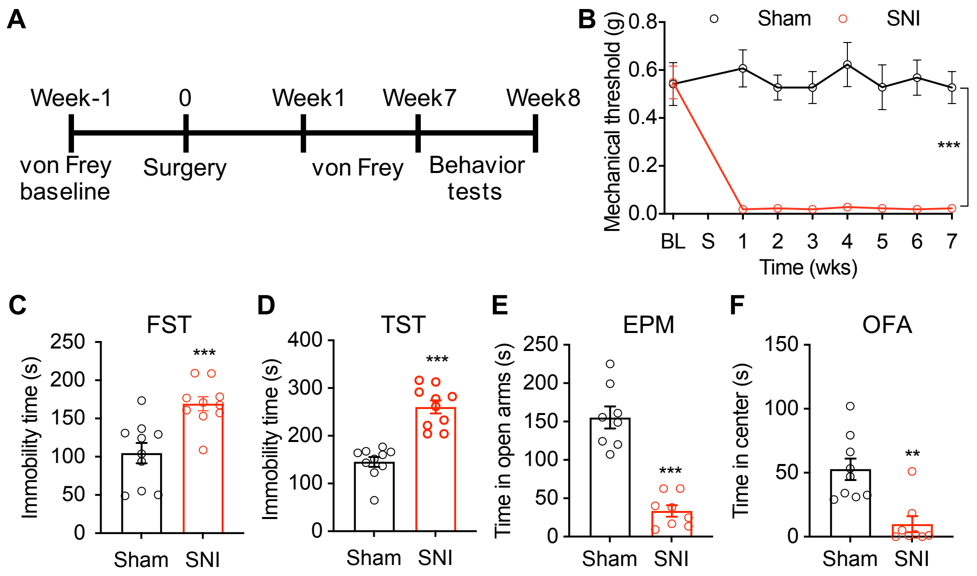


**Fig. S1. Depressive/anxiety-like behaviors are induced by chronic neuropathic pain.** (**A**) Experimental paradigm. (**B**) Time course of SNI-induced sensory pain (n = 7 mice for each group). BL, baseline; S, sham or SNI surgery. (**C-F**) Neuropathic pain mice (7 weeks after SNI surgery) displayed depressive/anxiety-like behaviors, as shown by increased immobile times in the forced swim test (FST) and the tail suspend test (TST) and reductions in the duration in open arms in the elevated plus maze (EPM) test and time in center in the open filed activity (OFA) test (n = 8-9 mice for each group). Data are means ± s.e.m. ** $P < 0.01$, ***$P < 0.001$. Two-way ANOVA followed by Tukey's *post-hoc* test (B), two-tailed unpaired Student's *t*-test (C-F).

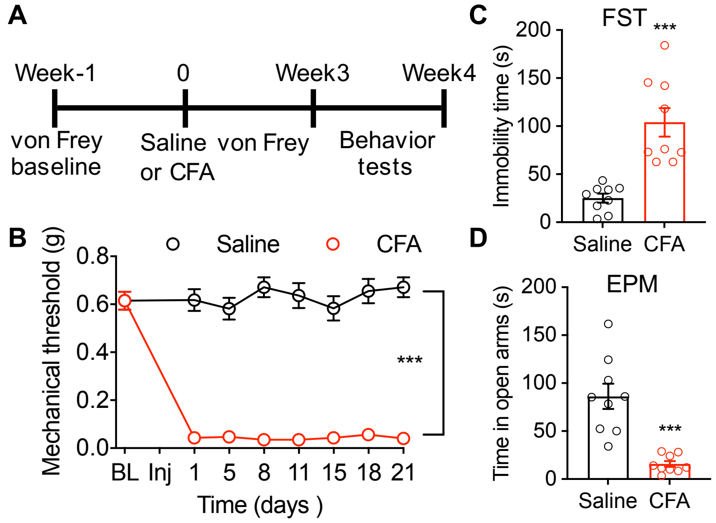


**Fig. S2. Depressive/anxiety-like behaviors are induced by chronic inflammatory pain.** (**A**) Experimental paradigm. (**B**) Time course of complete Freund's adjuvant (CFA)-induced sensory pain (n = 9 mice for each group). BL, baseline; Inj, saline or CFA injection. (**C** and **D**) Forced swim test (FST) and elevated plus maze (EPM) show that CFA treatment induced depressive/anxiety-like behaviors (n = 9 mice for each group). Data are means ± s.e.m. $***P < 0.001$. Two-way ANOVA followed by Tukey's *post-hoc* test (B), two-tailed unpaired Student's *t*-test (C and D).

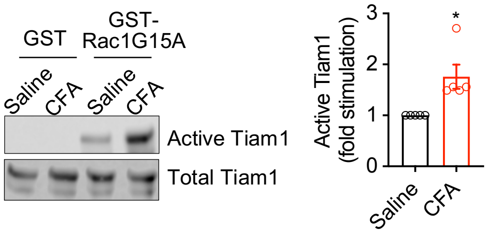


**Fig. S3. Chronic inflammatory pain activates Tiam1 in the ACC.** Active Tiam1 was detected by an affinity-precipitation assay using GST-Rac1G15A, which preferentially binds to activated GEFs. Tiam1 was precipitated from lysates prepared from the ACC of mice 3 weeks after saline or CFA injection. Total Tiam1 levels are also shown. (n = 5 mice for each group). Data are means ± s.e.m. * $P < 0.05$. Two-tailed unpaired Student's *t*-test.

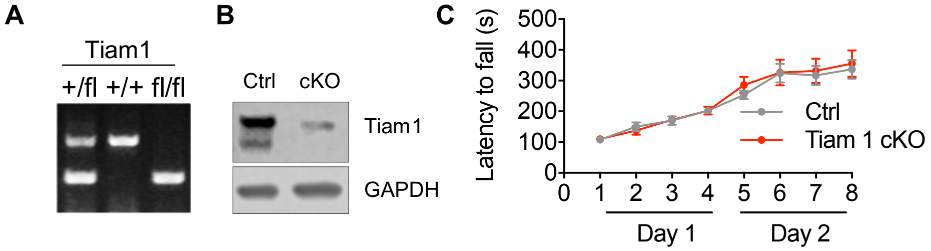


**Fig. S4. *Tiam1* cKO mice.** (**A**) PCR genotyping of tail DNA prepared from Tiam1$^{+/fl}$, Tiam1$^{+/+}$, and Tiam1$^{fl/fl}$ mice. (**B**) Representative immunoblots of forebrain lysate from control littermates (Ctrl) and *Tiam1* cKO (*Tiam1*$^{flox/flox}$::*CamKIIα*-Cre) mice probed with antibodies against Tiam1 and GAPDH. (**C**) Control (Ctrl) and *Tiam1* cKO mice were tested on an accelerating rotarod for two days (4 trials per day) and their motor performance was compared. No significant difference was detected between the two groups of mice (Ctrl, n = 12 mice; cKO n = 14 mice). Two-way ANOVA followed by Tukey's *post-hoc* test (C).

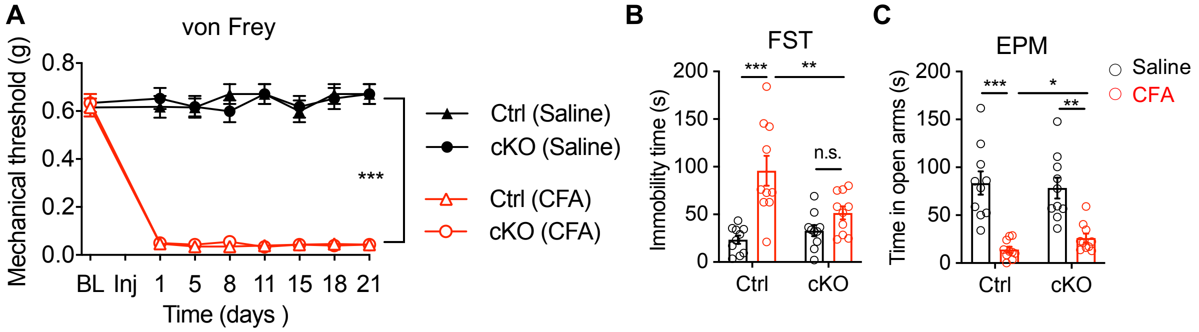


**Fig. S5. Genetic deletion of Tiam1 from postnatal forebrain excitatory neurons reduces inflammatory pain-induced depressive/anxiety-like behaviors.** (**A**) Time course of CFA-induced sensory pain showing that *Tiam1* cKO mice displayed no difference in mechanical allodynia compared to controls (Ctrl) before or during the 3 weeks following saline or CFA injection (n = 10 mice for each group). BL, baseline. Inj, saline or CFA injection. (**B** and **C**) Behavioral tests demonstrating that *Tiam1* cKO mice showed reduced inflammatory pain-induced depressive/anxiety-like behaviors in FST and EPM tests compared to that in control mice (Ctrl). (n = 10 mice for each group). Data are means ± s.e.m. * $P < 0.05$, ** $P < 0.01$, ***$P < 0.001$, n.s., no significance. Two-way ANOVA followed by Tukey's *post-hoc* test.

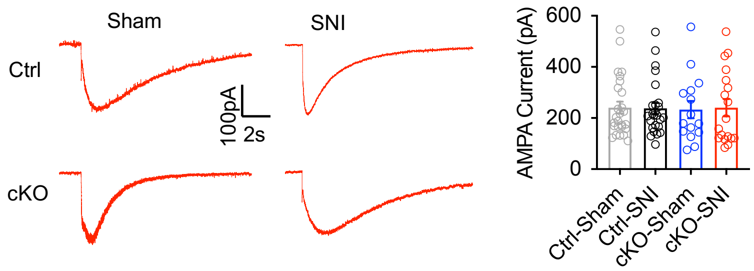


**Fig. S6. AMPAR currents elicited by puff application of AMPA.** Representative traces and mean changes in AMPAR currents elicited by puff application of 200 μM AMPA to ACC pyramid neurons in control (Ctrl) and *Tiam1* cKO mice 7 weeks after sham or SNI surgery (n = 16-26 neurons from 3 mice). Data are expressed as mean ± S s.e.m. One-way ANOVA followed by Tukey's *post hoc* test.

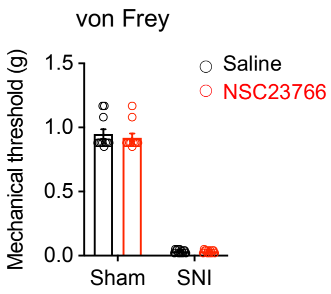


**Fig. S7. NSC23766 treatment (1 mg/kg) does not affect SNI-induced pain hypersensitivity.** Low dose of NSC23766 (1 mg/kg, i.p.) had no effect on SNI-induced pain hypersensitivity (7 weeks after SNI surgery). Data are means ± s.e.m. (n = 11 mice for each group). Two-way ANOVA followed by Tukey's *post-hoc* test.

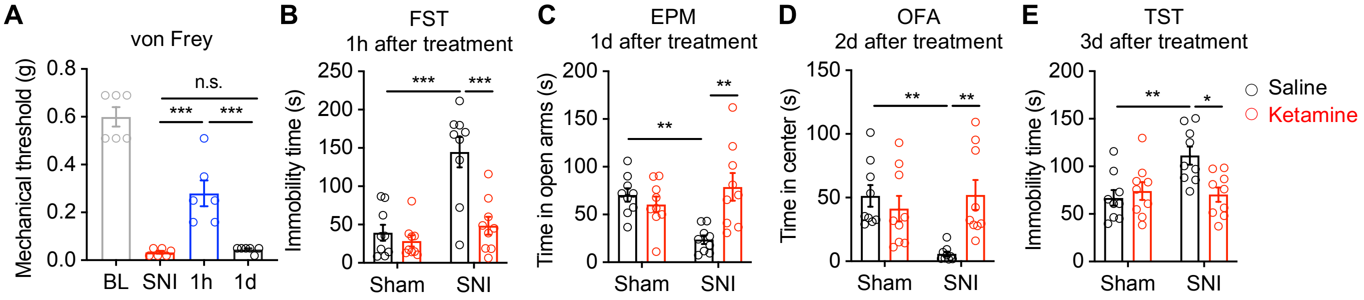


**Fig. S8. Antidepressant-like effect of a single systemic ketamine administration.** (**A**) A single ketamine dose relieved SNI-induced mechanical hypersensitivity at 1 hr, but not 1 day after administration (n = 6 mice). (**B**) FST showing a decrease in the immobility time of SNI mice 1 hr after ketamine injection (n = 9 mice for each group). (**C**) Ketamine treatment resulted in an increased time in open arms of SNI animals in the EPM test 1 day later (n = 9 mice for each group). (**D**) Ketamine treated SNI mice showed an increased time in center during OFA 2 days after administration (n = 9 mice for each group). (**E**) Ketamine reduced immobility times of SNI mice in the TST 3 days after administration (n = 9 mice for each group). Data are means ± s.e.m. * $P < 0.05$. ** $P < 0.01$. *** $P < 0.001$. n.s., no significance. One-way ANOVA followed by Tukey's *post-hoc* test (A), two-way ANOVA followed by Tukey's *post-hoc* test (B-E).

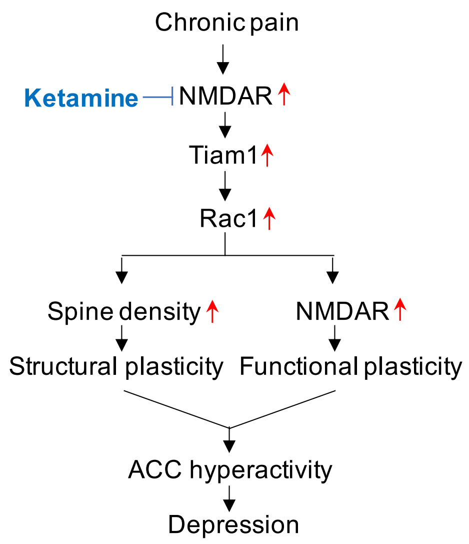


**Fig. S9. Proposed model.** Tiam1 links chronic pain-activated NMDARs to Rac1 activation in the ACC that orchestrates synaptic structural plasticity via spine remodeling and functional plasticity via synaptic NMDAR stabilization, which contributes to ACC hyperactivity and depression. Ketamine relieves depressive symptoms resulting from chronic pain by blocking Tiam1-mediated maladaptive plasticity in the ACC.
